# Haploinsufficiency of the essential gene RpS12 causes defects in erythropoiesis and hematopoietic stem cell maintenance

**DOI:** 10.1101/2021.05.04.442585

**Authors:** Virginia Folgado-Marco, Kristina Ames, Jacky Chuen, Kira Gritsman, Nicholas E. Baker

## Abstract

Ribosomal protein (Rp) gene haploinsufficiency can result in Diamond-Blackfan Anemia (DBA), characterized by defective erythropoiesis and skeletal defects. Some mouse Rp mutations recapitulate DBA phenotypes, although others lack erythropoietic or skeletal defects. We generated a conditional knockout mouse to partially delete *RpS12*, which results in homozygous embryonic lethality. *Rps12^+/-^* mice have growth and morphological defects, pancytopenia and impaired erythropoiesis. A striking reduction in hematopoietic stem cells (HSCs) and progenitors in the bone marrow (BM) was associated with decreased ability to repopulate the blood system after competitive and non-competitive BM transplantation. The mutants exhibited loss of HSC quiescence, which was associated with ERK and MTOR activation and increased global translation in HSC and progenitors. Thus, RpS12 has a very strong requirement in maintaining HSC quiescence and function, in addition to erythropoiesis that is affected in DBA patients.

## Introduction

In the cell, protein synthesis is one of the most energetically expensive processes, and both the specificity and overall level of translation are tightly regulated. The ribosome is the macromolecular machine tasked with translating mRNAs into proteins and, as such, plays an essential role in the physiology of the cell. Ribosomes are evolutionarily conserved ribonucleoprotein complexes composed of ribosomal RNA (rRNA) and ribosomal proteins (Rp) (Yonath and Franceschi 1998; Wilson and Doudna Cate 2012). They catalyze protein synthesis in all cell types, providing a supply line of steady-state levels of necessary cellular proteins (Wilson and Doudna Cate 2012). The functional components of the ribosome are highly conserved, and in higher eukaryotes consist of a small subunit (40S) and a large subunit (60S). These ribosomal subunits contain a total of 79 ribosomal proteins in eukaryotes, including 34 ribosomal proteins that are also conserved in prokaryotes (Petibon et al. 2020). In most cell types, ribosomal protein genes are among the most highly expressed genes (Geiger et al. 2012; Ji et al. 2019). Most ribosomal proteins are essential for ribosome biogenesis and function, which makes them essential for cell growth and proliferation (de la Cruz et al. 2015).

Given the importance of ribosomes, mutations in components of the ribosome or the ribosome biogenesis pathway in humans result in a group of diseases known as ribosomopathies. Despite the essential role of the ribosome in all cell types, this group of diseases is characterized by the presence of defects in specific tissues. Heterozygous loss of function mutations in Rp genes lead to Diamond-Blackfan Anemia (DBA), a congenital bone marrow failure syndrome characterized by macrocytic anemia, skeletal defects, and increased cancer risk. In DBA patients, mutations have been identified in 21 out of the 79 existing Rp genes, along with the GATA1 transcription factor (Ulirsch et al. 2018). Strikingly, in approximately 30-40% of DBA patient cases a mutation has not yet been identified. The fact that only a subset of all Rp genes have been found altered in DBA patients poses the question of whether mutations in any Rp gene can result in DBA and, if not, what would be the consequences for mutations in those Rp genes.

The generation and characterization of mice with mutations in Rp genes in recent years has begun to shed light on this question. Mutations in Rp genes of both the large and the small ribosomal subunits have been found to have similar phenotypes to those of DBA patients, such as impaired erythropoiesis, skeletal defects, and increased incidence of cancer, including some *Rp* not yet implicated in DBA (Oliver et al. 2004; McGowan et al. 2008; Jaako et al. 2011; Terzian et al. 2011; Morgado-Palacin et al. 2015; Schneider et al. 2016). However, erythropoietic defects are not always reported for mutants in Rp genes, including some that are implicated in human DBA (Matsson et al. 2004; Watkins-Chow et al. 2013; Kazerounian et al. 2016). In addition, a variety of other defects are reported in particular genotypes, ranging from embryonic lethality to brain defects, pigmentation defects, and defects in other aspects of hematopoiesis (McGowan et al. 2008; Kondrashov et al. 2011; Terzian et al. 2011; Watkins-Chow et al. 2013; Morgado-Palacin et al. 2015).

The *RpS12* gene, which is not yet implicated in DBA, reportedly has special functions in *Drosophila* that differ from those of most Rp. Heterozygous loss of 66 out of the 79 *Drosophila Rp* genes result in a ‘Minute’ phenotype, named for its small adult sensory bristles and also characterized by delayed development (Marygold et al. 2007). Additionally, ‘Minute’ *Rp^+/-^* cells are eliminated by wild-type (WT) neighboring cells when they are found together in developing tissues, by a process known as cell competition (Morata and Ripoll 1975; Clavería and Torres 2016; Baker 2020). Remarkably, delayed development, reduced translation, cell competition and other aspects of the ‘Minute’ phenotype depend on the RpS12 protein, which seems to be required for haploinsufficient effects of other *Rp* genes, suggesting that RpS12 acts as a sensor or reporter of deficits in other Rp. Accordingly, increasing the copy number of *RpS12* enhances these ‘Minute’ phenotypes caused by mutations in other *Rp* genes, whereas reducing the *RpS12* gene copy number suppresses them (Kale et al. 2018; Boulan et al. 2019; Ji et al. 2019). Interestingly, *RpS12* is one of the few Rp genes whose null mutation does not present a ‘Minute’ phenotype in heterozygosis (Marygold et al. 2007; Kale et al. 2018). In mammals, it has been shown that RpS12 deletions are frequent in diffuse large B cell lymphoma samples, and that RpS12 distribution in the ribosomes was altered under hypoxic conditions in the human embryonic kidney cell line, HEK293, resulting in changes of their translatome (Derenzini et al. 2019; Brumwell et al. 2020). Human RpS12 has also emerged as a candidate regulator of Wnt secretion in cancer cells (Katanaev et al. 2020). However, the phenotype of *RpS12* deletion in mammals has not been determined.

Protein synthesis regulation is important in stem cells. To maintain proper homeostasis, hematopoietic stem cells (HSCs) sustain the balance between a quiescent and an actively dividing state (Cabezas-Wallscheid et al. 2017). Quiescent HSCs require low rates of protein synthesis, and even HSCs exiting quiescence still exhibit significantly lower translation rates than in more differentiated progenitors. Both increases and decreases in protein synthesis levels can impair HSC function (Signer et al. 2014; Hidalgo San Jose et al. 2020).

The AKT/MTORC1 signaling pathway is one of the most well-known signaling pathways that regulates translation, in part through the expression of ribosomal proteins and translation factors (Fonseca et al. 2014). Hyperactivation of AKT signaling pathway is deleterious for normal HSC function, and results in increased HSC cycling, with depletion of the stem cell pool (Yilmaz et al. 2006; Kharas et al. 2010; Lee et al. 2010; Magee et al. 2012). Activation of AKT by stem cell factor (SCF) and other growth factors leads to the activation of MTOR, which results in the phosphorylation of the ribosomal protein S6 kinase 1 (S6K1) and the protein initiation factor 4E binding protein1 (4EBP1) (Gentilella et al. 2015). Phosphorylation of S6 by S6K1 at Serine 235/Ser236 is associated with increased protein translation (Krieg et al. 1988; Roux and Topisirovic 2018). Additionally, phosphorylation of 4E-BP1 by MTOR at the Thr37 and Thr46 residues primes it for dissociation of 4E-BP1 from eIF4E, also activating translation (Schalm et al. 2003). Furthermore, another important pathway regulating growth and translation, the MEK/ERK pathway, has been shown to phosphorylate S6 at Serine 235/Ser236 promoting the translation preinitiation complex in mammalian cells (Roux et al. 2007).

To further explore the specific functions of Rp genes, their potential involvement in DBA, and the regulation of translation, we determined the phenotype of RpS12 deletion in mice. We generated a conditional knock-out mouse, *RpS12^flox/flox^,* which, when crossed to embryonically expressed Ella-Cre recombinase, allowed us to generate homozygous (*RpS12^-/-^)* and heterozygous knock-out mice (*RpS12^+/-^*). We report that, while homozygous loss of *RpS12* is lethal at early stages of embryogenesis, the *RpS12^+/-^* phenotype includes reduced body size, skeletal defects, and, in some cases, hydrocephalus and stroke. Similar to DBA patients, and some other previously published Rp mouse mutants, *RpS12^+/-^* mice present a block in erythroid maturation, lower red cell counts, and decreased spleen size. However, loss of RpS12 also leads to a striking reduction in the number of hematopoietic stem cells in the bone marrow, as well as significantly altered progenitor populations, leading to overall reduced bone marrow cellularity and a decreased ability of *RpS12^+/-^* BM cells to repopulate the blood system, uncovering an impairment in HSC and progenitor function. These phenotypes were associated with increased translation and loss of quiescence in HSCs.

## Results

### RpS12 haploinsufficiency results in a pleiotropic phenotype, including delayed growth and increased mortality

To test the role of the RpS12 protein in a mammal, we used CRISPR gene editing to generate a mouse line with *LoxP* sites flanking exons 2 and 3 of the endogenous *RpS12 locus* (*RpS12^flox^*) (Supp Fig.1). Excision of these 2 exons generates an allele that cannot produce functional RpS12 protein, since exon 2 contains the ATG translation initiation codon. We chose not to eliminate the entire *RpS12* locus, to avoid deleting the small nucleolar RNA genes Snord100 and Snora33, which are located in introns 4 and 5, respectively (Supp Fig. 1). We crossed *RpS12^flox/flox^* mice to a line that expresses the Cre recombinase embryonically (*EIIa-Cre*) to obtain RpS12 heterozygous knock-out (KO) mice (*RpS12^KO/+^)* (Fig. 1A). Unlike heterozygous null flies, which don’t have any observable phenotype (Marygold et al. 2007; Kale et al. 2018), *RpS12^KO/+^* mice have reduced growth rates post-partum in comparison to their wild-type littermates (Fig. 1B, C). Additional phenotypes include kinked tails, mild hyperpigmentation of the footpads, and an increased risk of hydrocephalus (Fig. 1D, E, F). Although hydrocephalus has not been reported previously, some of these phenotypes have also been found in other ribosomal protein (Rp) mutant mouse models (Oliver et al. 2004; McGowan et al. 2008; Terzian et al. 2011). Furthermore, *RpS12^KO/+^* mice have increased mortality, especially in early post-natal stages, most of which is associated with hydrocephalus or the inability to gain weight (Fig. 1G).

**Figure 1.**
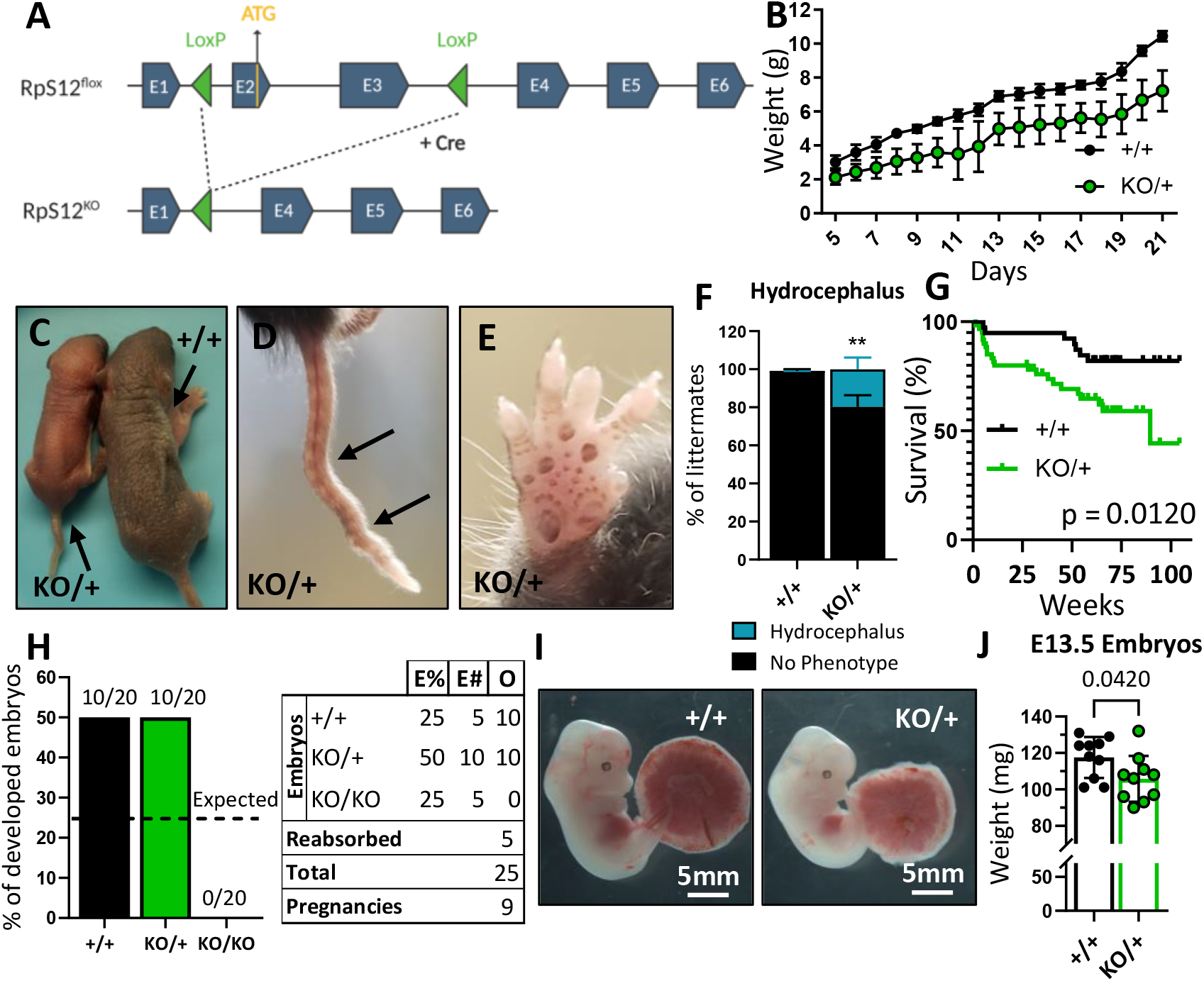
Loss of one copy of RpS12 results in delayed growth, morphologic defects, and reduced viability. **(A)** Conditional *RpS12^flox^* transgenic knock-in has two loxP sites flanking exons 2 and 3, that are removed by Cre recombinase activity to generate *RpS12^KO^*. **(B)** Post-natal growth curve of *RpS12^KO/+^* and *RpS12^+/+^* littermates (+/+ n=8 and KO/+ n=11 pups). **(C)** Picture of 5-day-old *RpS12^KO/+^* and *RpS12^+/+^* littermates. **(D)** Representative picture of “kinked” tail in *RpS12^KO/+^* mouse. **(E)** Representative picture of the anterior footpad hyperpigmentation in *RpS12^KO/+^*. **(F)** Quantification of the percentage of mice presenting hydrocephalus per litter (n=27 litters, 2-way ANOVA p=0.0035). **(G)** Kaplan-Meier survival curves of *RpS12^KO/+^* and RpS12^+/+^ littermates starting at day 5 of age (+/+ n=39 and KO/+ n=60, log-rank Mantel-Cox test p=0.012). **(H)** Embryo genotype segregation from crosses between RpS12^KO/+^ male and female. Graph represents percentage of developed embryos and the table shows the total numbers (E%=expected percentages, E#=expected numbers, O=observed numbers). **(I)** Representative pictures of E13.5 embryos with their placentas. **(J)** E13.5 embryo weights (n=10 on each genotype, unpaired t-test p=0.0420).

To investigate if the KO allele of *RpS12* is lethal in the homozygous state, and whether *RpS12^KO/+^* animals have reduced growth during embryonic development, we crossed heterozygous *RpS12^KO/+^* male and female mice and analyzed the resulting embryos at stage E13.5 (we could not assess frequencies in pups at birth because *RpS12^KO/+^* females invariably died during labor). There were no *RpS12^KO/KO^* specimens among the embryos obtained (Fig. 1H), which led us to conclude that this genotype must be lethal prior to stage E13.5. Furthermore, *RpS12^KO/+^* embryos are smaller in size compared to their wildtype counterparts (Fig. 1I, J). Therefore, these results indicate that *RpS12* is an essential gene, whose homozygous loss leads to early embryonic lethality, and heterozygous loss causes reduced growth starting in embryogenesis, in addition to other defects recognized post-partum.

### Heterozygous loss of RpS12 results in erythropoiesis defects that worsen with age

We sought to understand if, similar to other Rp mutant mouse models, RpS12 heterozygous mutants have anemia or defective erythropoiesis. Analysis of peripheral blood counts showed that “young” (6-8 weeks old) *RpS12^KO/+^* mice had lower number of white blood cells (WBC), red blood cells (RBC), and platelets, a condition known as pancytopenia (Fig. 2A). We also observed a high mean corpuscular volume (MCV), which is reminiscent of the macrocytic anemia seen in DBA patients.

**Figure 2.**
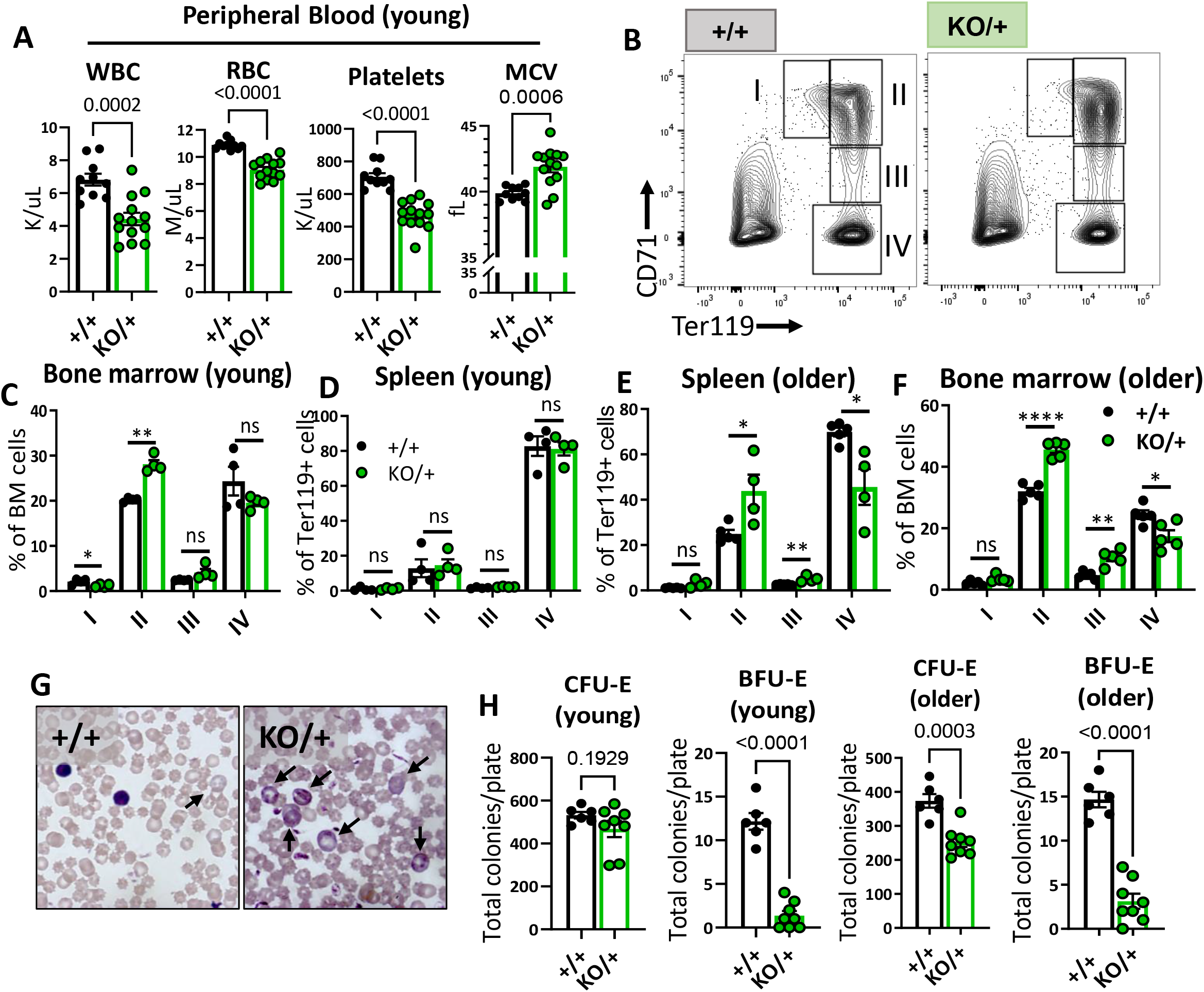
Haploinsufficiency of *RpS12* results in erythropoiesis defects that worsen with age. **(A)** Quantification of peripheral blood counts from young (6-8 weeks) littermates (+/+ n=10 and KO/+ n=13) (WBC=white blood cells, RBC=red blood cell, MCV=mean corpuscular volume). **(B)** Representative flow cytometry gating of bone marrow cells from 6-8-weeks-old mice of erythropoietic populations using Ter119 and CD71 markers. **(C, D, E, F)** Frequencies of erythroid progenitors in bone marrow and spleen of young (6-7 weeks old, +/+ n=4 and KO/+ n=4) and older (6-7 months old, +/+ n=5 and KO/+ n=5) mice. **(G)** Representative images of stained peripheral blood smears indicating the presence of reticulocytes (arrows). **(H)** Total number of CFU-E and BFU-E colonies per plate (5×10^5^ BM cells plated) in methylcellulose media supplemented with EPO (M3434) from young mice (6-7 weeks old, +/+ n=4 and KO/+ n=4, each biological sample had two replicates) and older mice (6-7 months old, +/+ n=4 and KO/+ n=4, each biological sample had two replicates). Statistical analysis: quantifications represent mean +/-SEM, when only two groups were being compared, unpaired t-test was performed, and for multiple comparison one-way ANOVA analysis was used. *p < 0.05, **p < 0.01, ***p < 0.001, ****p < 0.0001

To analyze erythropoiesis in *RpS12^KO/+^* mice, we used flow cytometry with the lineage markers Ter119 and CD71 on bone marrow and spleen cells (Fig. 2B). These populations represent different maturation stages of the red blood cell production process, which we refer to as RI (CD71^+^, Ter119^-^, proerythroblasts), RII (CD71^+^Ter119^+^, basophilic erythroblasts), RIII (CD71^mid^, Ter119^+^, late basophilic and polychromatophilic erythroblasts), and RIV (CD71^-^Ter119^+^, orthochromatic erythroblasts) (Socolovsky et al. 2001). Bone marrow samples from young *RpS12^KO/+^* mice showed a defective transition between the RII and RIII stage cells, while the erythropoiesis in spleen populations was unchanged (Fig. 2C, D). This impairment in erythropoiesis worsened with age, as samples from “older” (6-7 month-old) *RpS12^KO/+^* mice had a higher accumulation of RII and RIII stage cells, while RIV population numbers were decreased in both spleen and bone marrow samples at this age (Fig. 2E, F). In agreement with defective erythropoiesis, peripheral blood samples showed a visibly elevated percentage of reticulocytes in Wright-Giemsa stained blood smears (Fig. 2G). To assess erythropoietic progenitor function, we performed colony-forming unit (CFU) assays in methylcellulose media optimized for the differentiation of erythroid progenitors. Consistent with the observed impairment of erythropoiesis, *RpS12^KO/+^* bone marrow cells generated fewer BFU-E colonies, indicating reduced erythroid progenitor function (Fig. 2H). Altogether, these results show that RpS12 is required for erythroid differentiation, and demonstrate a role for Rps12 in erythropoiesis, similar to what has been observed in mouse models of DBA genes like *RpL11* or *RpS19* (Jaako et al. 2011; Morgado-Palacin et al. 2015).

### RpS12^KO/+^ mice have a striking reduction in hematopoietic progenitor populations, resulting in chronic pancytopenia

We were intrigued by the fact that *RpS12^KO/+^* mice have pancytopenia (Fig. 2A), since this is not a common feature of DBA patients. Due to the general decrease of peripheral blood cell numbers in *RpS12^KO/+^* mice, we hypothesized that hematopoietic stem and progenitor cells (HSPCs) might be affected. Using flow cytometry analysis, we assessed the stem cell and progenitor populations in the bone marrow using previously defined markers (Pietras et al. 2015) (Fig. 3A). Indeed, compared to the controls, *RpS12^KO/+^* revealed a striking reduction in the numbers of long-term HSCs (LT-HSCs: Flk2^-^CD48^-^CD150^+^ Lineage^-^Sca1^+^c-kit^+^ (LSK)) and short-term HSCs (ST-HSCs: Flk2^-^ CD48^-^CD150^-^ LSK) (Fig. 3B). In addition, in *RpS12^KO/+^* bone marrow, the numbers of all hematopoietic progenitor populations were significantly reduced (Fig. 3C). Accordingly, compared to the WT littermates, young (6-8-week-old) *RpS12^KO/+^* mice had lower bone marrow cellularity and decreased spleen weights (Fig.3D, E). Additionally, older (6-7-month-old) *RpS12^KO/+^* mice also had lower HSC numbers (Fig. 3F). Interestingly, we observed a partial recovery of some of the HSPC populations with age, such as multi-potent progenitors (MPP) 2 and 3, and the granulocyte-macrophage progenitors (GMP), as well as normalized overall BM cellularity, but not of spleen size (Fig. 3G, H, I). This, however, did not lead to improved blood counts (Fig. 3J), indicating that HSPC function was not significantly improved with age. Lastly, since RpS12 deletion resulted in decreased HSC and progenitor numbers, we assessed the self-renewal capacity of *RpS12^KO/+^* bone marrow cells. Plating assays in complete methylcellulose media showed a decreased clonogenic activity of *RpS12^KO/+^* bone marrow cells, as evidenced by the lower number of total colonies observed in the first round of plating (Fig. 3K). Additionally, *RpS12^KO/+^* cells have reduced serial replating capacity, suggesting decreased self-renewal capacity (Fig. 3K). Together, these results suggest that RpS12 plays an essential role in HSC function, including self-renewal and differentiation.

**Figure 3.**
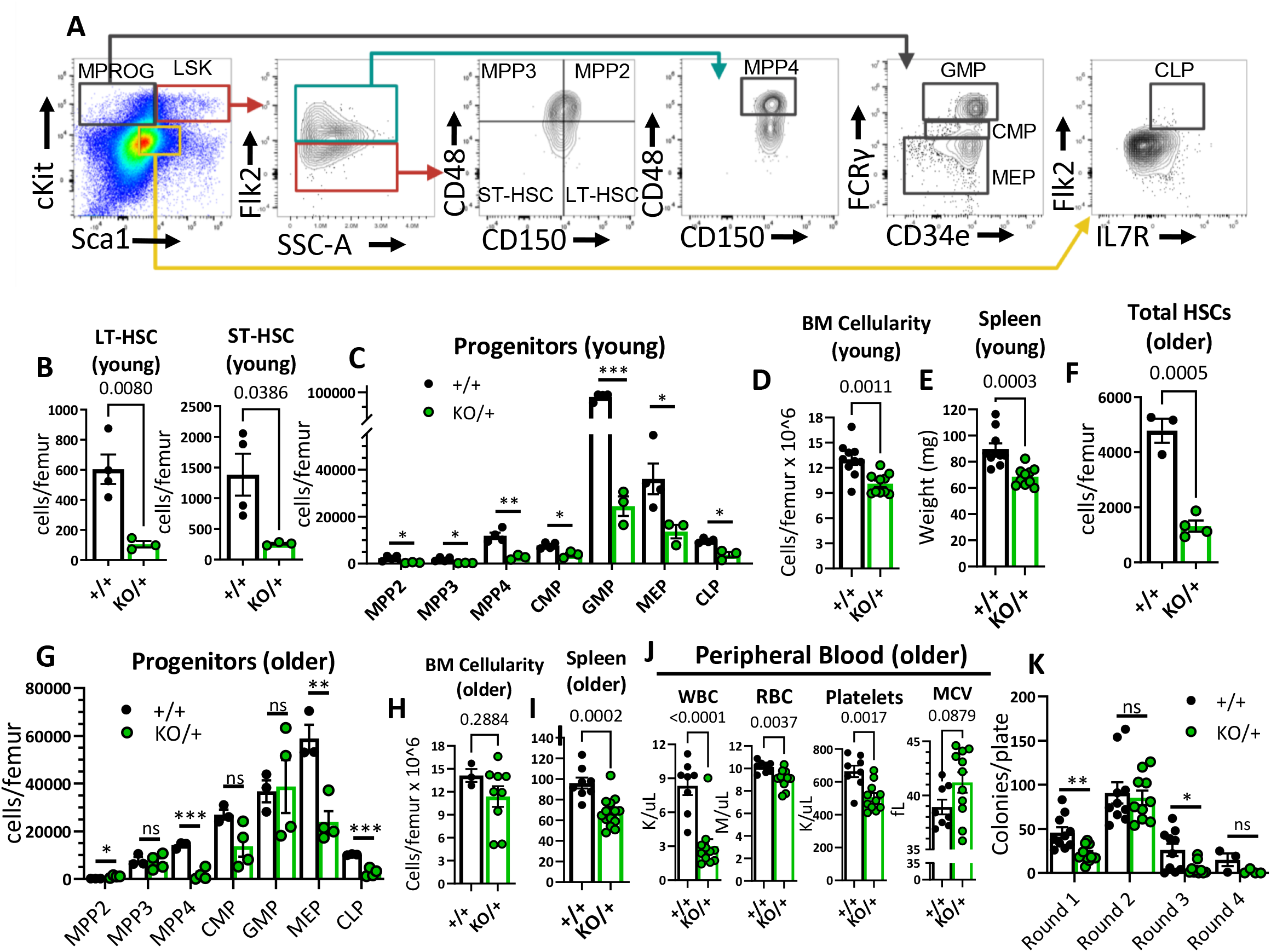
Reduced HSCs and other hematopoietic progenitor numbers in RpS12^KO/+^ mice. **(A)** Representative gating strategy used to identify bone marrow populations of LSKs: long-term HSC (LT- HSC), short-term HSC (ST-HSC), multi-potent progenitors (MPP2, MPP3, MPP4) and myeloid progenitors (MPROG): common myeloid progenitor (CMP), granulocyte-monocyte progenitor (GMP), megakaryocyte-erythrocyte (MEP) and common lymphoid progenitor (CLP). **(B)** Total LT-HSCs and ST-HSCs per femur of young mice (6-8 weeks old littermates, +/+ n=4 and KO/+ n=3). **(C)** Total number of cells per femur of indicated hematopoietic progenitor populations in young mice (6-8 weeks old littermates, +/+ n=4 and KO/+ n=3). **(D)** Bone marrow cellularity represented as cells per femur ×10^6^ from young mice (6-7 weeks old littermates, +/+ n=10 and KO/+ n=10). **(E)** Spleen weights of young (6-7 weeks old, +/+ n=10 and KO/+ n=10) mice. **(F)** Total HSCs per femur of older mice (6–7-month-old, +/+ n=3 and KO/+ n=4). **(G)** Total number of cells per femur of indicated hematopoietic progenitor populations in older mice (6–7-month-old, +/+ n=3 and KO+ n=4). **(H)** Bone marrow cellularity represented as cells per femur ×10^6^ from older mice (older: 6-7 months old, +/+ n=3 and KO/+ n=9). **(I)** Spleen weights of older (6-7 months old, +/+ n=8 and KO/+ n=14) mice. **(J)** Quantification of peripheral blood counts from older mice (6-7 months old, +/+ n=8 and KO/+ n=11). **(K)** Total number of colonies per plate (1×10^4^ BM cells from 6-7-month-old mice plated in round 1 and 1×10^4^ cells plated from previous plate on each re-plating round) on each round of re-plating in complete methylcellulose media (+/+ n=5 and KO/+ n=5, 2 replicates per biological sample). Statistical analysis: quantifications represent mean +/-SEM, unpaired t-tests were performed to established significance among populations between genotypes *p < 0.05, **p < 0.01, ***p < 0.001, ****p < 0.0001

### Heterozygous loss of RpS12 impairs ability of HSCs to reconstitute peripheral blood

We assessed self-renewal and differentiation properties of *RpS12^KO/+^* bone marrow cells (CD45.2+) *in vivo* by transplanting bone marrow into lethally irradiated B6.SJL mice (CD45.1+) (Fig. 4A). Interestingly, compared to the *RpS12^Flox/Flox^ or RpS12^Flox/+^* controls, *RpS12^KO/+^* bone marrow recipients had decreased survival, with 5 out of 20 transplanted mice dying within the first 8 weeks in the *RpS12^KO/+^* group vs 0 out of 20 dying in the control group (Fig. 4B). Whereas 100% of the control recipients were able to reconstitute the bone marrow in the long term (up to 20 weeks), only 55% of the *RpS12^KO/+^* recipients did so (Fig. 4C). Furthermore, longitudinal analysis of donor chimerism in the peripheral blood revealed that compared to the controls, surviving *RpS12^KO/+^* transplant recipients had significantly decreased donor chimerism (%CD45.2+) in the B and T cell lineages, and a trend toward decreased chimerism in the myeloid lineage (Fig. 4D-G). Together, this data suggests that bone marrow cells that lack RpS12 are deficient in hematopoietic repopulating capacity after lethal irradiation.

**Figure 4.**
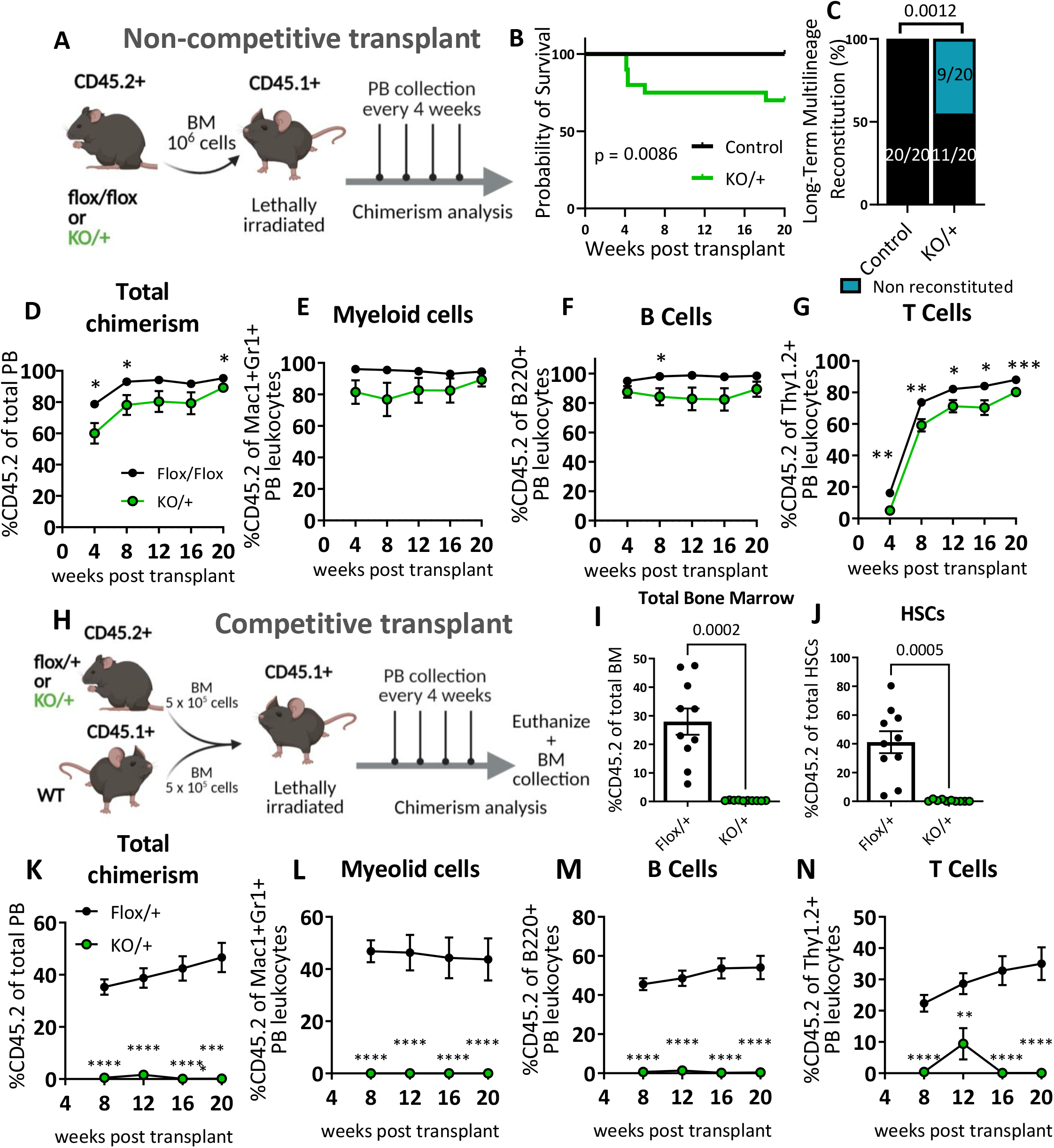
Heterozygous loss of *RpS12* impairs HSCs ability to reconstitute peripheral blood. **(A)** Non-competitive BM transplant strategy testing the long-term reconstituting activity of *RpS12^KO/+^* HSCs. 10^6^ bone marrow cells from *RpS12^KO/+^* or *RpS12^flox/flox^* samples (CD45.2+) were transplanted into lethally irradiated B6.SJL (CD45.1+) mice, peripheral blood chimerism was determined every 4 weeks. **(B)** Kaplan-Meier survival curves of mice transplanted with BM cells from *RpS12^KO/+^* and control *RpS12^flox/+^* or *RpS12^flox/flox^* mice (control n=20 and KO/+ n=20 transplanted mice, combination of 2 independent non-competitive transplants with 1 donor per genotype transplanted into 10 host mice each). **(C)** Frequency of recipient mice with long-term (20-weeks) multi-lineage reconstitution (≥0.5% in all three macrophages, B, and T cells)(control n=20 and KO/+ n=20 transplanted mice, combination of 2 independent non-competitive transplants). **(D-G)** Peripheral blood donor derived **(D)** total chimerism and **(E-G)** multi-lineage chimerism in non-competitively transplanted whole bone marrow (CD45.2+) recipients (flox/flox n=10 and KO/+ n=10). **(H)** Schematic representation of the competitive bone marrow transplant. 5×10^5^ cells from *RpS12^KO/+^* or *RpS12^flox/+^* donor bone marrow (CD45.2+) mixed with 5×10^5^ competitor bone marrow cells from B6.SJL (CD45.1+) mice were injected into lethally irradiated B6.SJL (CD45.1+) mice. Chimerism in peripheral blood was determined every 4 weeks and bone marrow chimerism was analyzed at 20 weeks after transplant. **(I)** Total bone marrow chimerism and **(J)** HSCs donor-derived (CD45.2+) chimerism in the recipient bone marrow (Flox/+ n=10 and KO/+ n=10 competitive-transplanted mice). **(K-N)** Donor derived peripheral blood chimerism of competitively transplanted *RpS12^KO/+^* or *RpS12^flox/+^* bone marrow cells as described in **H**. Non-competitive transplants were performed twice, using different controls: RpS12^flox/+^ or RpS12^flox/flox^. The competitive transplant was performed once, using RpS12^flox/+^ mice as a control group Statistical analysis: data represent mean +/-SEM, unpaired t-tests were performed to assess significance among populations between genotypes *p < 0.05, **p < 0.01, ***p < 0.001, ****p < 0.0001

To assess *RpS12^KO/+^* bone marrow cell repopulation capacity under more stringent conditions, we performed competitive transplantation of control (*RpS12^Flox/+^*) or *RpS12^KO/+^* bone marrow (CD45.2^+^) mixed with competitor WT bone marrow from B6.SJL mice (CD45.1^+^) in a 1:1 ratio into lethally irradiated B6.SJL (CD45.1^+^) recipient mice. Post-transplantation we monitored donor chimerism in the peripheral blood over time and analyzed the bone marrow chimerism at 20 weeks post-transplantation (Fig. 4H). Compared to the controls, *RpS12^KO/+^* transplant recipients showed a striking decrease in the percentage of donor-derived bone marrow cells and of HSCs (Fig. 4 I, J), accompanied by significant and persistent reduction in peripheral blood total donor chimerism in both myeloid and lymphoid lineages (Fig. 4K-N). Together, these data suggest that RpS12 haploinsufficiency leads to perturbed HSC self-renewal, resulting in ineffective hematopoiesis.

### The embryonic hematopoietic system is largely unaffected in *RpS12^KO/+^* animals

The striking reduction of hematopoietic progenitor numbers in *RpS12^KO/+^* adult bone marrow prompted us to investigate if this phenotype could be a consequence of defective HSC production in the fetal liver during embryogenesis. We analyzed fetal liver hematopoietic populations of E13.5 *RpS12^+/+^* and *RpS12^KO/+^* embryos because it has been shown that HSC numbers increase, and differentiation begins, between days 12 and 16 of embryogenesis in this organ (Sugiyama et al. 2011). First, we looked at the gross morphology and cellularity of the liver, neither of which were significantly different between the genotypes (Fig. 5A, B). Next, we assessed different stages of erythropoiesis using Ter119 and CD71 markers as previously described in fetal liver (Magee and Signer 2021). Most of the cells in population V were lost during staining, and therefore we did not include them in our analysis. Compared to the control embryos, *RpS12^KO/+^* embryos showed no apparent impairment in erythropoiesis in the fetal liver (Fig. 5C, D). Finally, we analyzed the distribution of HSPCs in E13.5 embryos. Overall, we did not observe any significant changes in the frequencies of LT-HSCs (CD48^-^CD150^+^ LSK), ST-HSCs (CD48^-^ CD150^-^ LSK), MPP (CD48^+^ LSK), CMP, GMP or MEP populations (Fig. 5E-H). These results show that partial loss of RpS12 does not affect embryonic hematopoiesis by E13.5. Therefore, the later HSPC deficiency is not a consequence of a defect in embryonic specification.

**Figure 5.**
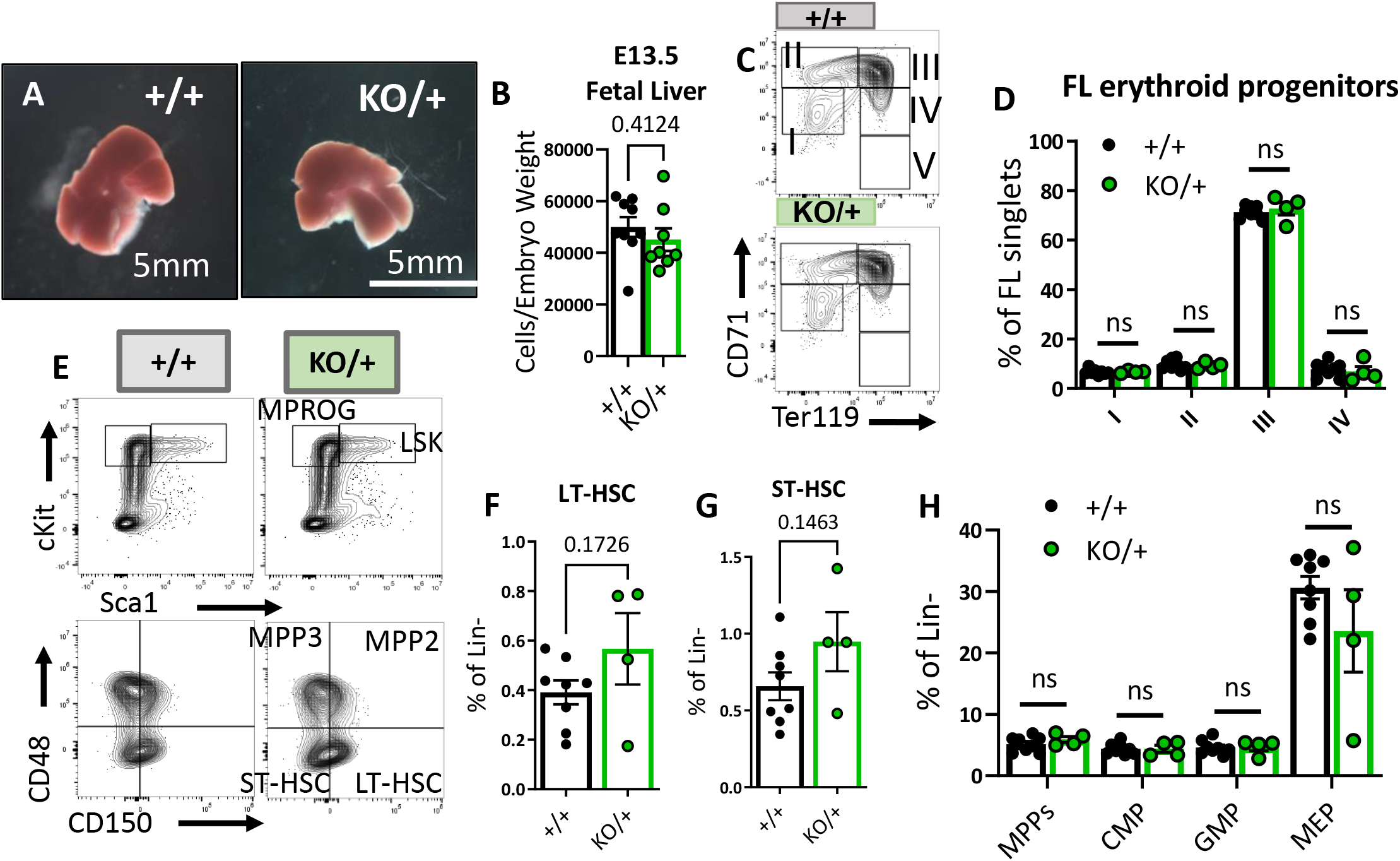
Embryonic hematopoietic system is largely unaffected in *RpS12^KO/+^* animals. **(A)** Representative images of RpS12^+/-^ and littermate E13.5 fetal livers. **(B)** Quantification of total number of cells per liver, normalized to embryo weight (+/+ n=9 and KO/+ n=8). **(C)** Representative flow cytometry gating of erythropoietic populations using Ter119 and CD71 markers of fetal liver samples from E13.5 embryos. (E) Representative flow cytometry gating of Lin- (top) and LSK (bottom) populations in E13.5 fetal livers. **(F,G,H)** LT-HSCs, ST-HSCs and indicated progenitor populations represented as percentages of the Lin- population in E13.5 fetal livers. **(D,F,G,H)** Biological samples are +/+ n=8 and KO/+ n=4. Statistical analysis: quantifications represent mean +/-SEM, unpaired t-tests were performed to established significance among populations between genotypes *p < 0.05, **p < 0.01, ***p < 0.001, ****p < 0.0001

### *RpS12^KO/+^* HSCs and some hematopoietic progenitors show higher translation, cycling and apoptosis

There are several important factors that maintain the HSC pool, including quiescence, low translation levels, and cell survival. We therefore analyzed the distribution of HSPCs among the cell cycle stages defined by the DNA content (Hoechst) and the levels of Ki67 (Fig. 6A). We observed a lower proportion of *RpS12^KO/+^* HSCs in the G0 stage of the cell cycle, and a significantly increased proportion in the actively cycling phases G1 and S/G2/M (Fig. 6B). Similar results were observed in MPP2/3 (LSK, Flk2^-^, CD48^+^), MPP4, and myeloid (MPROG) and common lymphoid progenitors (CLP) (Fig. 6B). These results show that compared to the control, *RpS12^KO/+^* HSCs are significantly less quiescent, with a higher proportion of HSCs and progenitors actively cycling.

**Figure 6.**
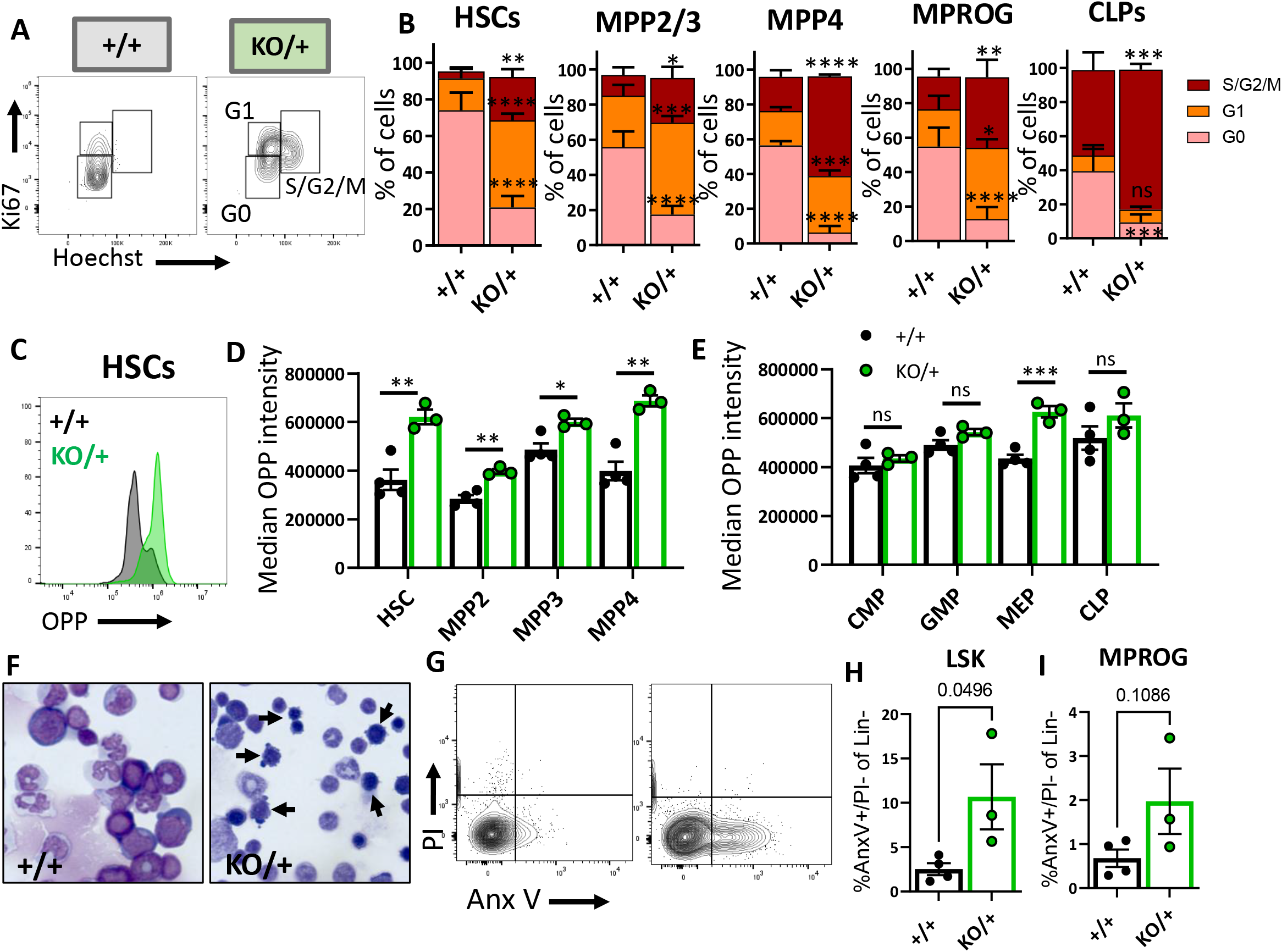
HSCs and other hematopoietic progenitors have altered cycling, global translation levels and apoptosis in *RpS12^KO/+^* bone marrow. **(A)** Representative flow cytometry gating of HSCs (Flk2^-^CD48^-^LSK) cell cycle stages (G0, G1, S/G2/M) distribution determined by DNA (Hoechst) and Ki67 levels. **(B)** Cell cycle stages distribution in HSCs and in indicated progenitor populations. Asterisks correspond to p values assessing significant differences in each cell cycle stage between *RpS12^KO/+^* and *RpS12^+/+^* mice (6-8-weeks-old littermates, +/+ n=4 and KO/+ n=3). **(C)** Representative flow cytometry histogram showing OPP intensity in *RpS12^KO/+^* (green) and *RpS12^+/+^* (grey) HSCs. **(D, E)** Median OPP intensity of the indicated bone marrow populations (6-8-weeks-old littermates, +/+ n=4 and KO/+ n=3). This analysis was repeated in 6-7-month-old mice with similar results. **(F)** Representative images of bone marrow cytospins showing high number of apoptotic cells (arrows) in *RpS12^KO/+^* samples. **(G)** Representative flow cytometry gating of LIN- population showing apoptotic populations as determined by AnexinV and PI staining. **(H, I)** Percentage of apoptotic (AnnexinV+) cells in LSK (Lin^-^cKit^+^Sca1^+^) and Myeloid progenitor (MPROG; Lin^-^cKit^+^Sca1^-^) populations (6-8-weeks-old littermates, +/+ n=4 and KO/+ n=3). Statistical analysis: quantifications represent mean+/-SEM, two-way ANOVA (B-F) and unpaired t-tests were performed to established significance among populations between genotypes *p < 0.05, **p < 0.01, ***p < 0.001, ****p < 0.0001

Cell cycle activation generally requires translation, but previous studies have reported a generalized decrease in global translation in some Rp mutants, including in HSPC, despite a decrease in HSC quiescence (Oliver et al. 2004; Signer et al. 2014; Schneider et al. 2016). To test the global translation levels of each HSPC population in RpS12 heterozygous mice using flow cytometry, we performed an *in vitro* assay on freshly isolated HSCs and progenitors using the puromycin analog o-propargyl puromycin (OPP) as previously described (Signer et al. 2014). Unexpectedly, compared to the *RpS12^+/+^* controls, *RpS12^KO/+^* HSCs and multipotent progenitor cell populations all showed increased levels of global translation (Fig. 6C, D). The difference was especially remarkable in HSCs. Interestingly, compared to the controls, *RpS12^KO/+^* myeloid progenitors did not exhibit differences in OPP intensity, and among different myeloid progenitor populations, only the megakaryocyte-erythrocyte progenitors (MEP) had a significant increase in OPP incorporation (Fig. 6E). Thus, these data suggest that a decrease in Rps12 leads to an abnormal increase in global protein translation in immature bone marrow populations, including HSCs.

Cell death can deplete the HSC pool, and can result from chronic HSC activation. We asked whether this reduction of HSCs in *RpS12^KO/+^* animals is due to an increase in apoptosis. Interestingly, compared to controls, *RpS12^KO/+^* animals have an increased number of apoptotic cells in bone marrow cytospins (Fig. 6F). To quantify the level of apoptosis in the immunophenotypic populations of the bone marrow cells, we used the flow cytometry markers PI and Annexin V together with population-specific cell surface markers (Fig. 6G). Our flow analysis confirmed a significant increase in apoptosis in Lineage^-^Sca1^+^c-Kit^+^ (LSK) cells, a population that contains HSCs and MPPs, but not in more mature myeloid progenitors (Fig. 6H, I).

### *RpS12^KO/+^* HSPCs have overactivated MEK/ERK and AKT/TOR signaling pathways

Because *RpS12^KO/+^* mutants have increased translation, we assessed the activity of the AKT/MTOR pathway, since it is known to regulate translation. Since the AKT/MTOR pathway is activated by stem cell factor (SCF), we determined the level of the AKT/MTOR pathway activation in the presence and absence of SCF, by assessing the phosphorylation levels of phospho-AKT (Ser 473) and the MTOR downstream effectors phospho-S6 (Ser235/236) and phospho-4E-BP1 (Thr37/46). Our results show that, in the more immature LSK population, which is enriched for HSCs and MPPs, the levels of p-AKT, p-S6 and p-4E-BP1 were significantly elevated in *RpS12^KO/+^* animals compared to wild-type littermates, not only upon stem cell factor (SCF) stimulation, but even at the non-stimulated baseline. Indeed, the *RpS12^KO/+^* genotype alone was a more potent activator than SCF (Fig. 7A-C). Interestingly, this was not the case for the more mature myeloid progenitor cells, where the levels of p-AKT, p-S6 and p-4E-BP1 are comparable to the controls in both non-stimulated and SCF-stimulated conditions, and these cells also exhibited more normal translation rates (Fig. 7D-F). Since phosphorylation of S6 and 4EBP1 leads to increased translation, this data corroborates the increase in translation observed in the *RpS12^KO/+^* LSK population (Fig. 6D). Interestingly, more mature MPROG do not have increased translation (Fig. 6E) and do not have increased activation of the AKT/MTOR pathway (Fig. 7D-F). Together, these data suggest that activation of translation and the increase in AKT/MTOR signaling in *RpS12^KO/+^* mutant cells are specific to HSCs and MPPs.

**Figure 7.**
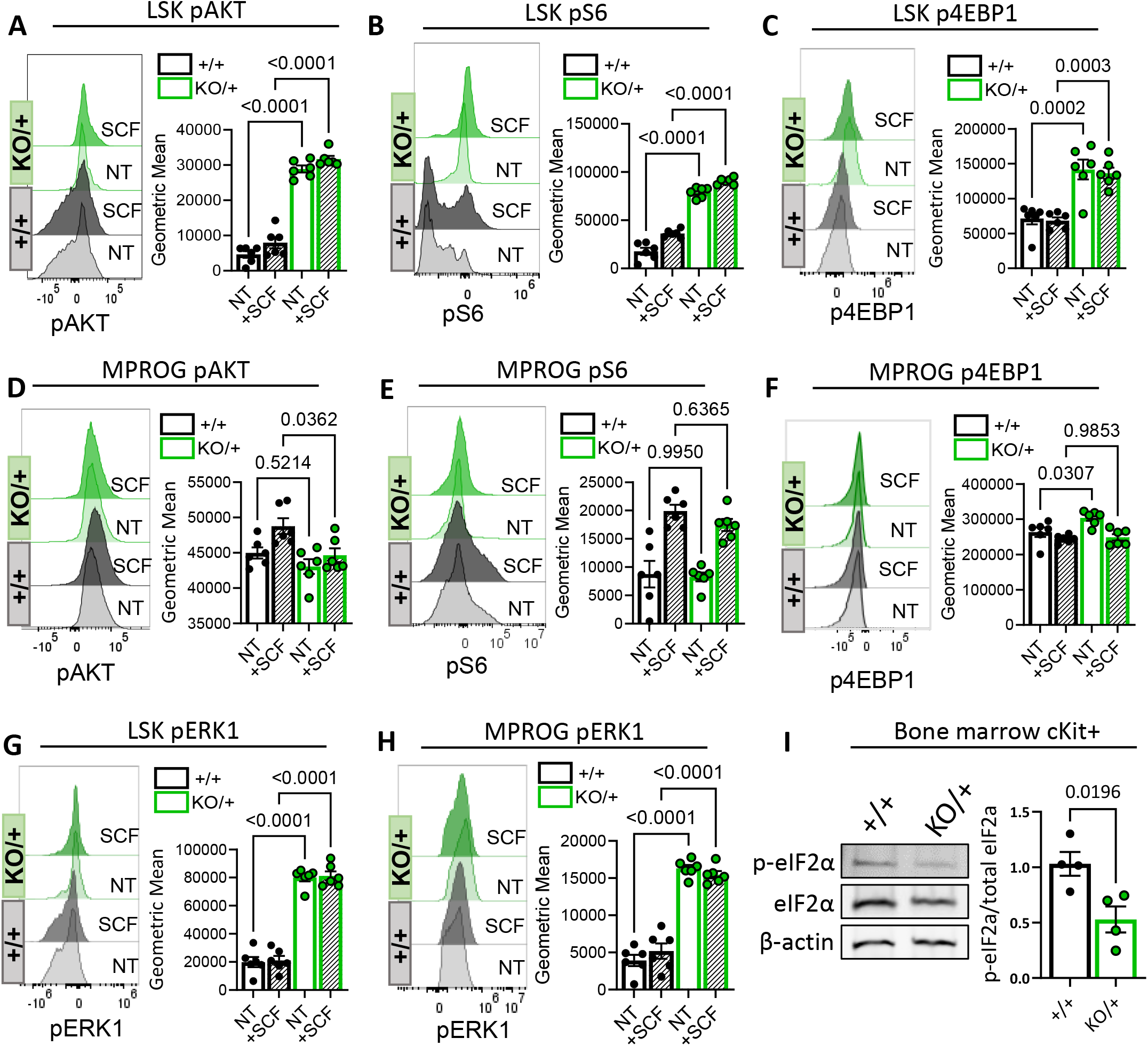
Decreased Rps12 levels leads to the excessive activation of the AKT/MTOR and ERK signaling pathways. **(A-H)** Representative phospho-flow cytometry histograms and quantification of the normalized geometric mean fluorescent intensity of pAKT (Ser 473) **(A,D)**, pS6 (Ser235/236) **(B,E)**, p4EBP1 (Thr37/46) **(C,F)** and pERK1(Thr202/Tyr204) **(D,H)** signal in the LSK **(A,B,C,G)** and MPROG **(D,E,F,H)** bone marrow cell populations. Baseline signal was determined in the none treated (NT) serum starved cells, stimulation was done with the stem cell factor (SCF) *ex vivo* for 5 minutes. Immunophenotypic populations were defined as follows: <u>LSK:</u> Lin^-^cKit^+^Sca1^+^, <u>MPROG:</u> Lin^-^cKit^+^Sca1^-^. **(I)** Representative images of western blot analysis and quantification of phospho-eIF2α normalized to the total eIF2α protein in cKit-enriched BM samples (6-8-weeks-old littermates, +/+ n=4 and KO/+ n=4) **(A-H)** 7-weeks-old littermates, +/+ n=6 and KO/+ n=6 biological samples were used. Statistical analysis: quantifications represent mean+/-SEM, one-way ANOVA Tukey’s multiple comparison’s test were performed to established significance among samples between genotypes

Additionally, compared to wild-type controls, *RpS12^KO/+^* mutant animals also have increased phospho-ERK1 (Thr202/Tyr204) in the LSK and MPROG populations under SCF stimulated and non-stimulated conditions (Fig. 7G, H). Another regulator of translation is the eukaryotic initiation factor 2α (eIF2α), which is required for CAP-dependent translation initiation. Cells respond to several stress conditions by phosphorylating eIF2α, reducing global translation and upregulating stress-response genes (Wek et al. 2006; Sigurdsson and Miharada 2018). In skeletal muscle stem cells, eIF2α phosphorylation promotes quiescence and stem cell maintenance (Zismanov et al. 2016). Interestingly, compared to the control cells, *RpS12^KO/+^* cKit+ bone marrow progenitor cells show decreased levels of p-eIF2α (Fig.7I), which also correlates with increased translation.

## Discussion

We generated an RpS12 knock-out mouse and describe the homozygous and heterozygous mutant phenotypes. Homozygous loss of *RpS12* was lethal during early embryogenesis. Similar to other *Rp* mutant mice, *RpS12* heterozygous mutants have reduced body size, skeletal defects, and anemia, and we showed that RpS12 is required for erythroid differentiation. Some of the mice also exhibited hydrocephalus. Heterozygous mice were viable with several visible phenotypes and blood cell defects. In many respects, *RpS12* mutant phenotypes resemble those of mice mutant for other *Rp* genes, including defective erythropoiesis, suggesting that *RpS12* could be a candidate gene for DBA.

Most strikingly, we report that RpS12 is also crucial for normal hematopoietic stem cell self-renewal and differentiation, with defective engraftment and long-term repopulation of *Rps12^KO/+^* bone marrow in transplantation experiments. This seems specific to adult HSCs, as no hematopoietic defect was observed in the fetal livers of *RpS12^KO/+^* embryos. Although we have not determined whether the transition from the fetal liver to the bone marrow occurs normally, such a defect would only be expected to delay bone marrow engraftment, and does not seem sufficient to explain the chronically defective HSC function and striking loss of HSC quiescence that we observe in *RpS12^KO/+^* adult mice.

The loss of bone marrow HSC quiescence was associated with chronic activation of Akt/mTor and Erk, increased translation, and HSC apoptosis. Chronic activation of the AKT/MTOR pathway has been shown to result in increased HSC cycling, apoptosis, and decreased self-renewal (Chen et al. 2008; Kharas et al. 2010), which could explain the HSPC exhaustion phenotype and increased HSPC apoptosis in *RpS12^KO/+^* mice. Fetal liver HSCs, which were unaffected in *RpS12^KO/+^*, normally have a higher proliferative activity and higher translation rates than adult HSCs (Magee and Signer 2021), which may be why they are less sensitive to heterozygous deletion of RpS12.

The increase in HSPC-specific, global translation upon heterozygous deletion of *RpS12* is the opposite of what has been reported in *RpL24^Bst/+^* and *RpS14^+/-^* HSPCs (Signer et al. 2014; Schneider et al. 2016), although there are other *Rp* genotypes where HSC cycling is increased (Terzian et al. 2011; Schneider et al. 2016). However, it is consistent with the increased translation that has been reported in mice with deletion of *Pten* in HSCs, which leads to activation of the AKT/MTOR pathway (Signer et al. 2014). Importantly, detailed studies of RpS12 depletion from ribosomes in yeast clearly demonstrate a strong reduction in translation as a result (Martin-Villanueva et al. 2020). Unless RpS12 has a different function in mice, or in HSPCs, this raises the possibility that enhanced translation in *RpS12^KO/+^* cells might be due, not to enhanced translation by ribosomes lacking RpS12, but to increased activity of the remaining intact ribosomes that contain RpS12. Such an increase would in fact be expected as a result of the striking activation of the AKT/MTOR and ERK pathways. *RpS12^KO/+^* c-Kit+ hematopoietic progenitors also have lower phosphorylated eIF2α, which would also predict higher translation levels.

It remains to be determined how *RpS12* haploinsufficiency leads to activation of the AKT/MTOR and ERK signaling pathways, which has not been reported in other *Rp* mutants. It is interesting that a regulatory role has been suggested for RpS12 in *Drosophila*, where multiple properties of *Rp* mutant cells, including translation, depend on RpS12-dependent activation of a transcriptional stress response mediated by the *Drosophila* transcription factor Xrp1, although the molecular mechanism is not yet known (Kale et al. 2018; Boulan et al. 2019; Ji et al. 2019). It is possible that mouse RpS12, either through an effect on specific translation or otherwise, plays a particular role in regulating HSC quiescence, through the direct or indirect activation of the AKT/MTOR and ERK pathways. However, it is also possible that HSC activation occurs indirectly, as a consequence of HSC apoptosis. In addition, we cannot exclude the possibility that RpS12 deletion in the bone marrow niche could lead to a loss of HSC quiescence through a non-autonomous mechanism. Specific RpS12 functions that have been suggested in mammalian cells and cancers could now be explored using this conditional knock-out model with tissue specific Cre-drivers (Derenzini et al. 2019; Brumwell et al. 2020; Katanaev et al. 2020).

The fact that *RpS12^KO/+^* mice exhibit fully-penetrant pancytopenia with a severe bone marrow failure phenotype, in addition to the erythropoiesis defect, raises the possibility that *RpS12* might not have been found mutated in DBA patients due to a more severe human phenotype that is not classified as DBA. Perhaps only a hypomorphic *RpS12* genotype could be associated with DBA. It is also the case, however, that caution is required extrapolating from mouse phenotypes to human. Nevertheless, our study raises the possibility that Rps12 could in fact be a candidate gene not only for DBA, but for a broader group of bone marrow failure disorders. One of the reasons that genetic alterations in RpS12 have not yet been reported in DBA or other bone marrow failure disorders could be that RpS12 is not included in the most common targeted next generation sequencing (NGS) panels used in the diagnosis of these disorders, some of which report molecular diagnostic rates of only 44-59% (Ghemlas et al. 2015; Muramatsu et al. 2017; Galvez et al. 2021). We suggest that in the future RpS12 should be included in expanded NGS panels for bone marrow failure disorders, or should be sequenced in the patients in whom there are no molecular findings in the standard NGS panels.

## Methods

### Generation of RpS12^flox^ knock-in mice

A pair of guide-RNAs (gRNAs) targeting intron 1 and intron 3 of *RpS12* gene respectively were designed by an online tool (http://crispr.mit.edu/) and generated by in vitro transcription. Cas9 mRNA was purchased from SBI. An *RpS12* conditional knockout homology-directed repair (HDR) plasmid containing 2kb homologous arms at each side and exon 2 and 3 flanked by loxP sites (Supp. Fig. 1) was generated by SLiCE cloning. Super ovulated female C57BL/6J mice (3–4 weeks old) were mated to C57BL/6J males, and fertilized embryos were collected from oviducts. The gRNAs, Cas9 mRNA and conditional knockout HDR plasmid were microinjected into the cytoplasm of fertilized eggs. The injected zygotes were transferred into pseudo pregnant CD1 females and the resulting pups were genotyped. Out of 20 pups, 2 mice were identified as *RpS12^flox/+^*, which were then crossed to obtain *RpS12^flox/flox^* (*Rps12^em1Nbakr^* MGI:6388411). The corresponding DNA sequences can be found in Supplementary table 1.

### Mice

All animals were housed at the Animal Housing and Studies Facility at Albert Einstein College of Medicine (AECOM) under pathogen-free conditions and experiments were performed following protocols approved by the Institutional Animal Care and Use Committee (IACUC) (Protocol #20181206). C57BL/6J and EIIa-Cre (FVB/N-Tg(EIIa-cre)C5379Lmgd/J) mice were obtained from Jackson. B6.SJL-Ptprca/BoyAiTac (CD45.1) mice from Taconic were used for transplantation experiments. To generate *RpS12^KO/+^* mice, we crossed *RpS12^flox/flox^* to EIIa-Cre mice and used primers flanking the floxed region to identify progeny where recombination had occurred. Unless indicated, these mice were kept as heterozygous by crossing them with C57BL/6J and the presence of the EIIa-Cre locus was crossed out. In each case, the genotypes were confirmed by PCR using genomic DNA extracted from tails using DNeasy kit from Qiagen (#69504). Peripheral blood samples were collected via facial vein bleeding under isoflurane anesthesia, and blood counts obtained using the Genesis analyzer (Oxford Science). To generate growth curves, 5-day-old pups were genotyped and numbered by cutting toes. Pups’ weight was measured daily from day 5 to day 21 of age.

### Preparation of the single cell suspension

Bone marrow single cell suspension was prepared from freshly harvested femurs, tibiae, ilia, and vertebrae by gentle crushing of the bones in phosphate buffered saline containing 2% fetal bovine serum (PBS/2% FBS) followed by filtration through a 40-µm strainer. Spleen cells were obtained from freshly harvested spleens. Single cell suspension was prepared by dissociating spleens using the flat end of a plunger against in 40-µm strainers and washed with PBS/2% FBS. Fetal liver single cell suspension was prepared from freshly harvested E13.5 fetal livers by passing through a 200-µl pipet tip and filtered through a 40-µm strainer in PBS/2% FBS. Cells were subjected to RBC lysis (Qiagen) according to the manufacturers protocol and used for the further steps described below. To calculate absolute number of cells per femur one femur per mouse was flushed, RBC lysed and cells counted to obtain total number of cells per femur.

### Flow cytometry on live cells

#### Bone marrow

Single cell bone marrow suspensions were stained with a cocktail of biotin-conjugated lineage antibodies for 30min at 4°C, washed with PBS/2% FBS, stained with fluorochrome-conjugated antibody cocktails for 30min at 4°C, washed with PBS/2% FBS, resuspended in PBS/2% FBS, filtered through a 40-µm strainer, and subjected to Flow analysis. For the antibody panels, refer to Supplementary Tables 2 and 3.

#### Peripheral blood

samples were subjected to RBC lysis, blocked with CD16/CD32 10min at 4°C followed by staining with fluorochrome-conjugated antibodies, washed with PBS/2% FBS, resuspended in PBS/2% FBS, filtered through a 40-µm strainer and subjected to Flow analysis. For the antibody panels, refer to Supplementary Table 2 and 3.

#### Erythropoiesis analysis

Obtained single cell suspension of spleen cells (without RBC lysis) was blocked with CD16/CD32 for 10 min at 4°C and stained with fluorochrome-conjugated antibodies for 30 min at 4°C (Supplementary Table 3). E13.5 fetal livers single cell suspension was blocked with CD16/CD32 10min at 4°C and stained with erythropoiesis or progenitor panels as described above. All blocking and staining steps were performed in PBS/2% FBS.

#### Apoptosis analysis

Single cell suspension samples were stained with the lineage-cocktail and fluorochrome-conjugated antibody cocktails as described above. After completion surface antibody staining, samples were incubated with FITC-conjugated Annexin V (BD-560931) and Propidium Iodide (BD-556463) following the manufacturer’s instructions.

All flow cytometry was performed with BD FACS LSRII or Cytek Aurora and data analysis was done with FlowJo Software (v9, v10).

### Flow cytometry on fixed cells

#### Cell cycle

Fresh single cell suspension of the RBC lysed bone marrow cells was stained with lineage antibodies followed by staining with fluorochrome-conjugated antibodies against surface markers, as described above. Immediately after staining cells were fixed and permeabilized using Cytofix/Cytoperm™ Fixation/Permeabilization Solution Kit (BD biosciences, BDB554714) according to the manufacturer’s instructions. Followed fixation, cells were incubated overnight at 4°C with FITC-conjugated Ki67 antibody in Perm/Wash buffer. DNA was stained with 25µl/ml Hoechst in Perm/Wash buffer before flow cytometry analysis. Flow cytometry was performed with BD FACS LSRII or Cytek Aurora and data analysis was done with FlowJo Software (v9, v10).

#### Global translation *in vitro*

protocol was based on a previously described assay (Signer et al. 2014). Single cell bone marrow RBC lysed cell suspension was obtained as described. Cells were resuspended in DMEM (Corning 10-013-CV) media supplemented with 50 µM β-mercaptoethanol (Sigma) and 20 µM OPP (Thermo Scientific C10456). Cells were incubated for 45 minutes at 37°C and then washed with Ca^2+^ and Mg^2+^ free PBS. The samples were stained with biotin-labeled antibodies, followed by staining with fluorochrome-conjugated antibody cocktails, fixed and permeabilized using Cytofix/Cytoperm™ as described above. After permeabilization with Perm/Wash buffer, cells were resuspended in Click-iT® Plus Reaction Cocktail (Thermo Scientific C10456) containing azide conjugated to Alexa Fluor 488 for 30 minutes at room temperature, washed once with Click-iT® Reaction Rinse Buffer and resuspended in Perm/Wash buffer. Flow cytometry was performed with Cytek Aurora and data analysis was done with FlowJo Software (v9, v10).

#### Phospho-flow cytometry

Bone marrow cells were starved for 1 hour in IMDM 2% FBS at 37°C, stained with lineage antibodies, followed by staining with fluorochrome-conjugated antibodies against surface markers, as described above. Post staining cells stimulated with 100ng/ml mSCF (Peprotech #250-03) in 2% PBS-FBS for 5 min at 37°C. Stained and stimulated cells were fixed and permeabilized with Cytofix/Cytoperm as described above and stained with phospho-S6 (Ser235/236) - Alexa 488 (Cell Signaling Technology, 4803S) (1:100) and phospho-AKT (Ser473) - Alexa647 (Cell Signaling Technology, 2337S), phospho-4E-BP1 (Thr37/46) -Alexa Fluor647 -(Cell Signaling Technology, 5123S) of pERK1(T202/Y204) - Alexa 488 (Cell Signaling 4374) at 1:20 dilutions. Cells were washed with Perm/Wash buffer to remove residual and unbound antibody, and resuspended in fresh Perm/Wash buffer, followed by flow cytometry analysis on the Cytek Aurora. Analysis of all flow cytometry data was performed using FlowJo software (v9, v10).

### Methylcellulose cultures and serial re-plating

Single cell bone marrow suspensions (post RBC lysis) or single cell fetal liver cell suspensions (without RBC lysis) were resuspended in RPMI media supplemented with 10%FBS and 1% penicillin/streptomycin. Cells were manually counted on a hemocytometer using Trypan blue and plated in methylcellulose media (M3434 or M3334, Stem Cell Technologies) at a density of 5×10^5^ live cells/ml (in M3434) or 10^4^ live cells/ml (in M3334) in 35mm cell culture plates. Samples were incubated at 37°C in 6.5% CO_2_ at constant humidity. Colonies were scored and evaluated 7-10 days after plating. To replate, cells were washed from the plates with RMPI media, counted, and re-plated in fresh M3334 methylcellulose at a density of 10^4^ live cells/dish in 35mm plates. This process was repeated until cell exhaustion in one of the experimental groups.

### Bone marrow transplantation

6-8-week-old B6.SJL (CD45.1) recipient mice were lethally irradiated with a single dose of 950 Gy using a Cesium-137 gamma-ray irradiator (Mark I irradiator Model 68) at least 3 hours before transplantation. For non-competitive assays, 10^6^ whole bone marrow cells from a donor control (*RpS12^flox/flox^* or *RpS12^flox/+^*) or *RpS12^KO/+^* mouse (CD45.2) were injected into the retro-orbital venous sinus of recipient mice under isoflurane anesthesia. For competitive transplants, 10^5^ whole bone marrow cells from control *RpS12^flox/+^* or *RpS12^KO/+^* donor mouse (CD45.2), and 10^5^ competitor cells from a B6.SJL (CD45.1) mouse were injected into each recipient mouse. Mice were given drinking water treated with 100mg/ml Baytril100 (Bayer) for 3 weeks after transplantation. Peripheral blood was collected every 4 weeks and animals were euthanized at the specified experimental time points.

### Western blot analysis

Whole bone marrow cells were enriched for cKit+ cells using CD117 MicroBeads and MACS LS Columns (Miltenyi Biotec 130-091-224 and 130-042-401) following the manufacturer’s protocol. 2 x 10^6^ cKit enriched cells were resuspended in 150 µl Laemmli buffer (BioRad 1610737) supplemented with 1:10 β-mercaptoethanol, passed through a 25-G needle to break the DNA and incubated at 95°C for 5 minutes. An equal amount of each sample was separated in polyacrylamide gels (BioRad 4568081), transferred to nitrocellulose membrane (Licor) and blocked (Licor 927-90001). IRDye near-infrared secondary antibodies (Licor) were used to visualize the proteins. The following primary antibodies were used: RpS12 (Proteintech; polyclonal), β-actin (Cell Signaling; 13E5), eIF2α (Cell signaling, 5324T), phospho-eIF2α (Thermo Scientific; Ser52; polyclonal).

### Histology

Peripheral blood smears and cytospins from RBC lysed bone marrow samples were stained using the Hema 3 System (Fisher) following the manufacturer’s instructions. The images were acquired using a Zeiss Axiovert microscope with a digital camera.

### Statistical methods

Two-tailed Student’s t-tests were performed to compare statistical significance between two samples. When comparing more than 2 groups, one-way ANOVA tests were performed with the Turkey’s multiple comparison test. For presence/absence of phenotype, statistical significance was calculated with Fisher’s exact test. Analysis was done using GraphPad Prism v9.

### Reagents

All antibodies used for flow cytometry assays, and primers used for CRISPR gene editing and PCR can be found in Supplementary Tables 1, 2, and 3. All other reagents are mentioned in the methods section.

## Acknowledgments

This work was supported by National Institutes of Health grants R01GM104213 (to N.E.B.), R01CA196973 (to K.G.), an award from the Albert Einstein College of Medicine Human Genetics Program (to N.E.B.), startup funds from the Albert Einstein College of Medicine and Albert Einstein Cancer Center (to K.G.), the NHLBI/NIH Ruth L. Kirschstein National Research Service Award F32HL146119 (to K.A.), and the IRACDA/BETTR training Institutional Research and Academic Career Development Award 2K12GH102779-07A1 (to K.A). For flow cytometry this work utilized the analyzers Cytek Aurora Multiparameter Flow Cytometer and BD LSR-II with the help from Dr. Jinghang Zhang, Dr. Yu Zhang and Aodengtuya Fnu. The Cytek Aurora Multiparameter Flow Cytometer was purchased with funding from the National Institutes of Health SIG grant #1S10OD026833-01. Data in this paper are from a thesis to be submitted in partial fulfillment of the requirements for the Degree of Doctor of Philosophy in the Biomedical Sciences, Albert Einstein College of Medicine.

K.G. receives research funding from iOnctura, S.A. The authors have no additional financial interests.

---

**Supplementary Figure 1.**
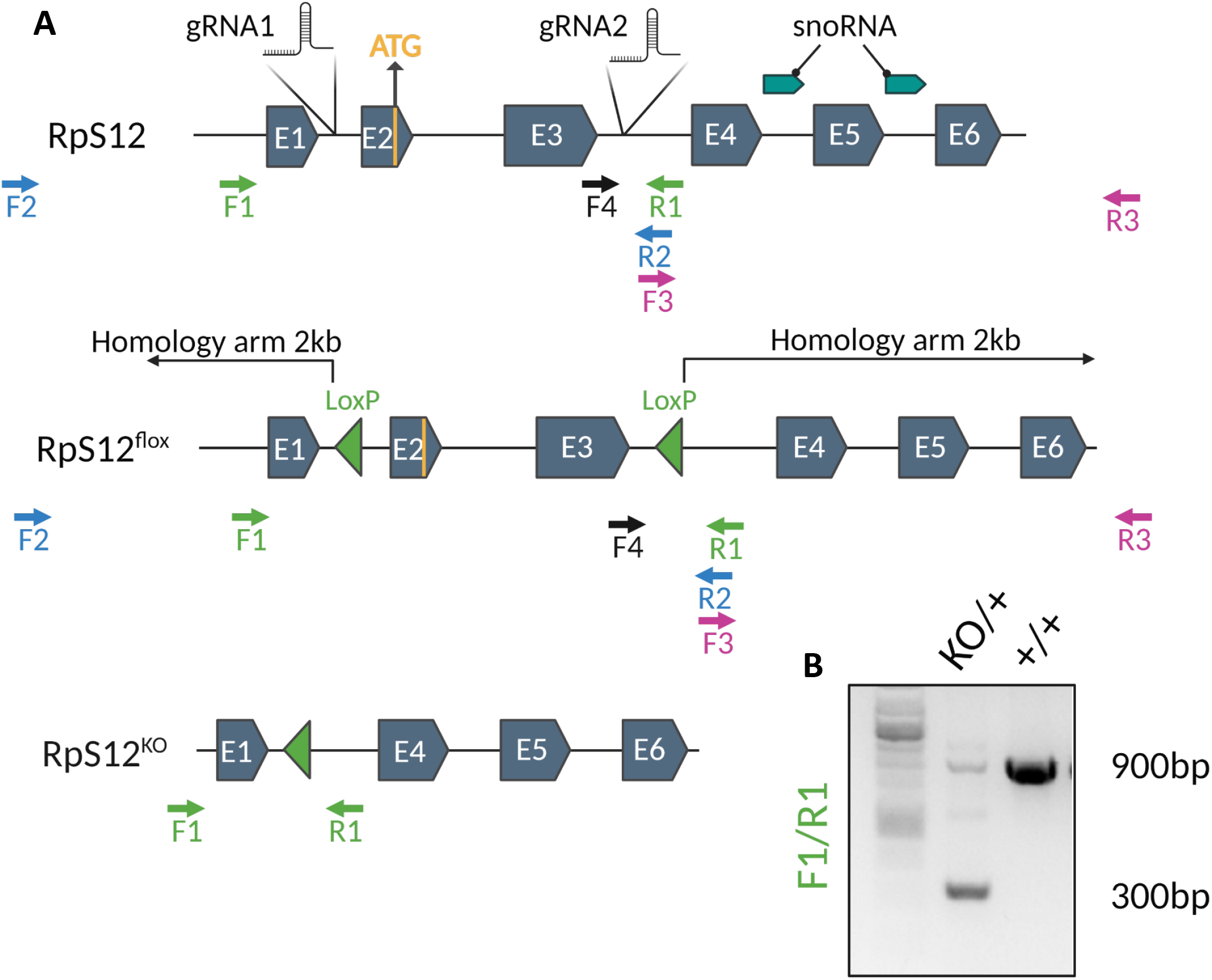
CRISPR gene editing and genotyping strategy for the generation of *RpS12^Flox^* and *RpS12^KO^*. **(A)** Diagram of the WT, Flox and KO alleles of RpS12 generated in this study indicating the position of Snord100 and Snora33 (snoRNA),Cas9 gRNAs target locations, and primers used for genotyping. The homology arms starting sites are indicated and the ends fall outside of the RpS12 locus. To identify the first transformants, two pair of primers were used for PCR amplification: F2/R2 and F3/R3. F2 and R3 fall outside of the sequence covered by the homology arms, to ensure the inserts are on the correct location. The presence of LoxP sites was confirmed by Sanger sequencing using primers F1 and F4 for F2/R2 fragments, and with F3 and R3 for F3/R3 fragments. To determine excision of exon 2 and 3 by Cre recombination primers F1 and R1 were used, which generate a 900bp fragment in *RpS12^+^* and a 300bp fragment in *RpS12^KO^* **(B)**.

**Supplementary table 1.**
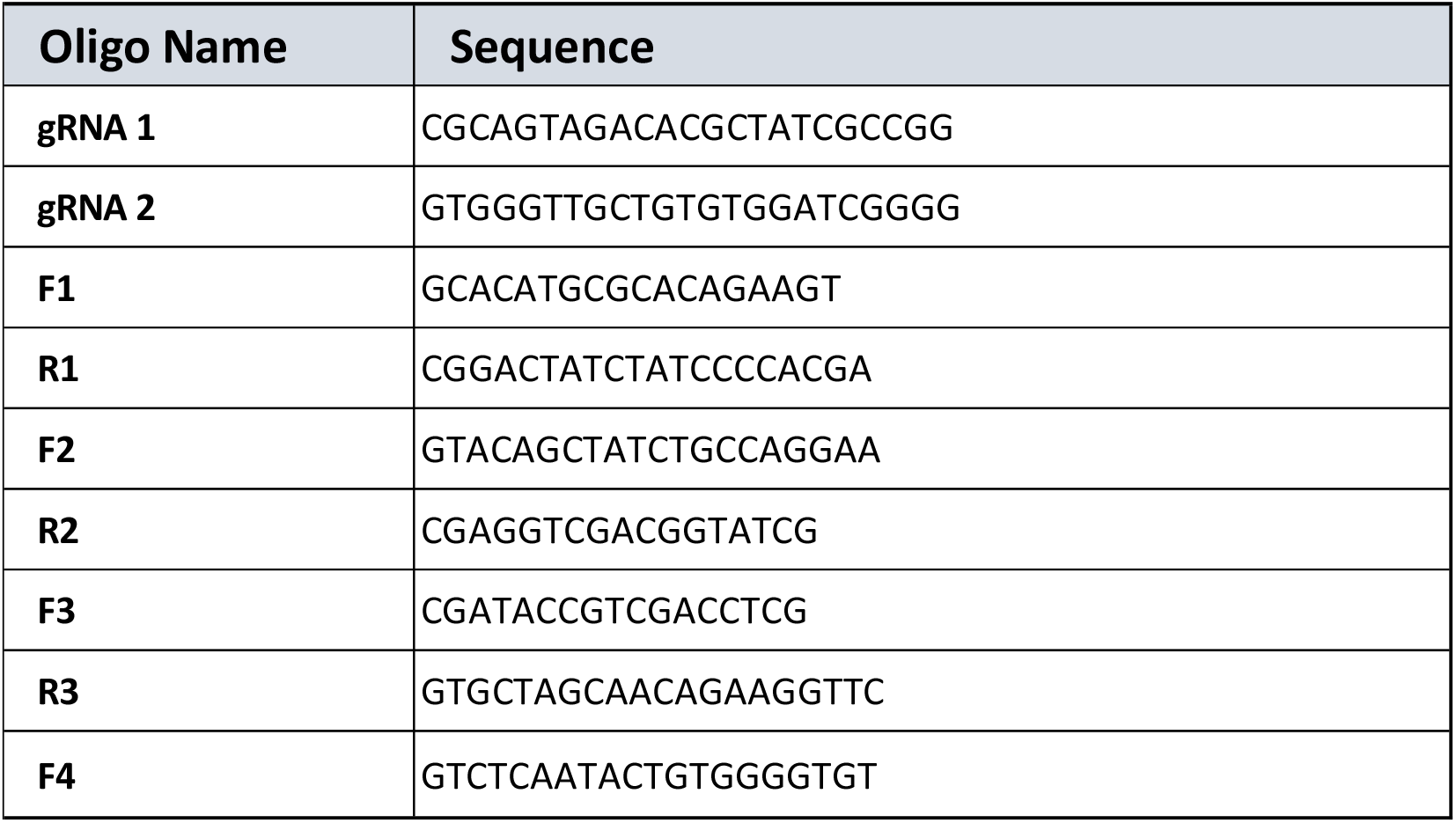

**Supplementary Table 2.**
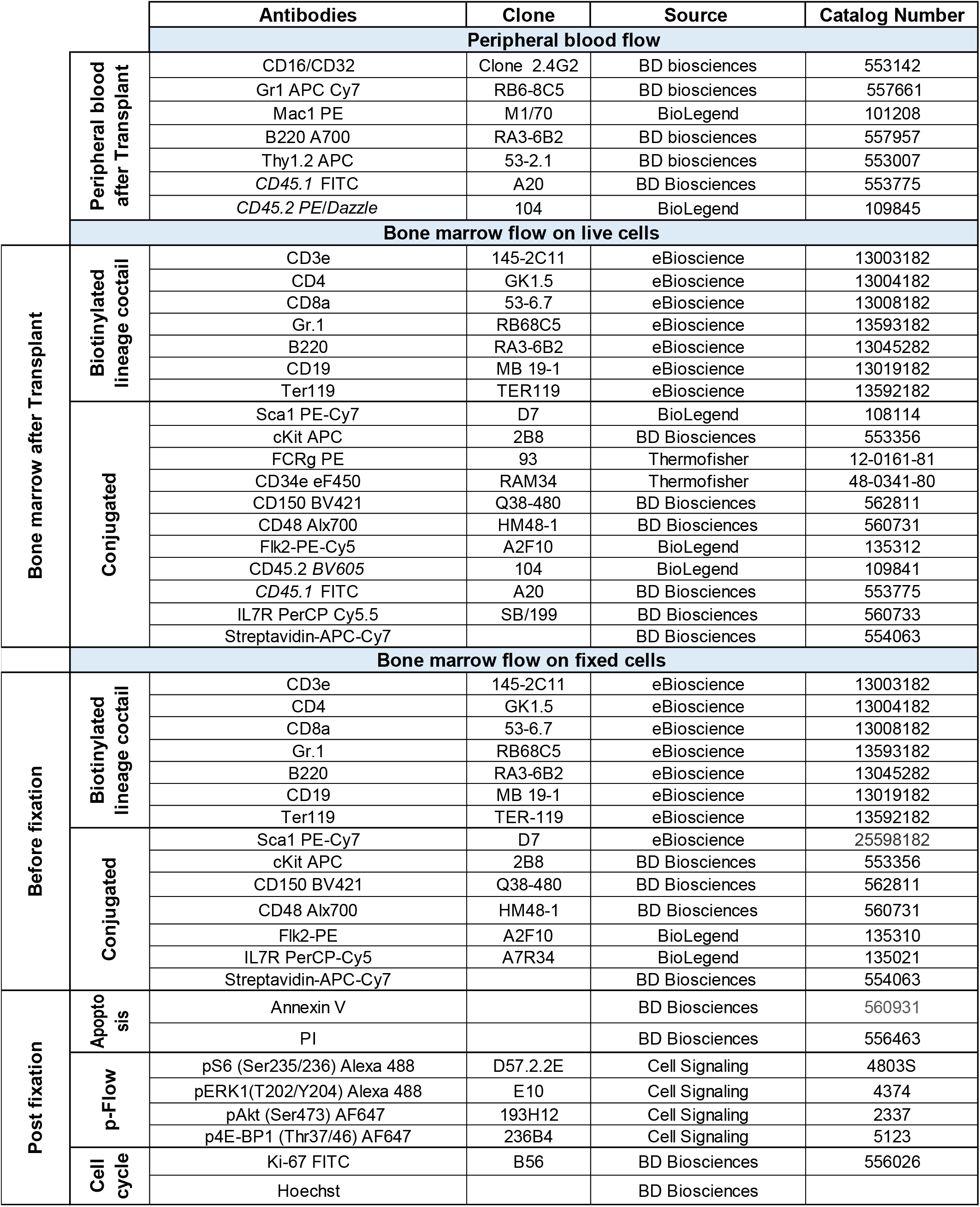

**Supplementary Table 3.**

|  |  | Antibodies | Clone | Source | Catalog Number |
| --- | --- | --- | --- | --- | --- |
| **Before fixation** | **Bone marrow flow OPP** |  |  |  |  |
|  | **Biotinylated lineage cocktail** | CD3e | 145-2C11 | eBioscience | 13003182 |
|  |  | CD4 | GK1.5 | eBioscience | 13004182 |
|  |  | CD8a | 53-6.7 | eBioscience | 13008182 |
|  |  | Gr.1 | RB68C5 | eBioscience | 13593182 |
|  |  | B220 | RA3-6B2 | eBioscience | 13045282 |
|  |  | CD19 | MB 19-1 | eBioscience | 13019182 |
|  |  | Ter119 | TER-119 | eBioscience | 13592182 |
|  |  | IL7R | eBioRDR5 | eBioscience | 13127882 |
|  | **Conjugated** | Sca1 PE-Cy7 | D7 | eBioscience | 25598182 |
|  |  | cKit APC | 2B8 | BD Biosciences | 553356 |
|  |  | FCRg PE | 93 | Thermofisher | 12-0161-81 |
|  |  | CD34e eF450 | RAM34 | Thermofisher | 48-0341-80 |
|  |  | CD150 BV421 | Q38-480 | BD Biosciences | 562811 |
|  |  | CD48 Alx700 | HM48-1 | BD Biosciences | 560731 |
|  |  | Flk2-PE Cy5 | A2F10 | BioLegend | 135311 |
|  |  | IL7R PerCP-Cy5 | A7R34 | BioLegend | 135021 |
|  |  | Streptavidin-APC-Cy7 |  | BD Biosciences | 554063 |
|  | **Non Conjugated** | OPP |  | Thermofisher | C10456 |
| **After** |  | Click-IT 488 |  | Thermofisher | C10456 |
|  |  | **Bone marrow and spleen (non-lysed)** |  |  |  |
|  |  | CD16/CD32 | Clone 2.4G2 | BD biosciences | 553142 |
|  |  | Ter119 APC | TER119 | BioLegend | 116211 |
|  |  | CD71 PE | C2 | BD biosciences | 561937 |

## Notes

### Competing Interest Statement

The authors have declared no competing interest.

